# How predictive can be predictions in the neurocognitive processing of auditory and audiovisual speech? A deep learning study

**DOI:** 10.1101/471581

**Authors:** Thomas Hueber, Eric Tatulli, Laurent Girin, Jean-luc Schwartz

**Affiliations:** Univ. Grenoble Alpes, CNRS, Grenoble INP, GIPSA-lab, 38000 Grenoble, France; Inria Grenoble Rhône-Alpes, 38330 Montbonnot, France

## Abstract

Sensory processing is increasingly conceived in a predictive framework in which neurons would constantly process the error signal resulting from the comparison of expected and observed stimuli. Surprisingly, few data exist on the amount of predictions that can be computed in real sensory scenes. Here, we focus on the sensory processing of auditory and audiovisual speech. We propose a set of computational models based on artificial neural networks (mixing deep feed-forward and convolutional networks) which are trained to predict future audio observations from 25 ms to 250 ms past audio or audiovisual observations (i.e. including lip movements). Experiments are conducted on the multispeaker NTCD-TIMIT audiovisual speech database. Predictions are efficient in a short temporal range (25-50 ms), predicting 40 to 60 % of the variance of the incoming stimulus, which could result in potentially saving up to 2/3 of the processing power. Then they quickly decrease to vanish after 100 ms. Adding information on the lips slightly improves predictions, with a 5 to 10 % increase in explained variance.

Interestingly the visual gain vanishes more slowly, and the gain is maximum for a delay of 75 ms between image and predicted sound.

## Introduction

The concept of “predictive brain” has progressively emerged in neurosciences in the 50s [1, 2]. It assumes that the brain is constantly exploiting the redundancy and regularities of the perceived information, hence reducing the amount of processing by focusing on what is new and eliminating what is already known. After half a century of experimental developments, the predictive brain has been mathematically encapsulated by Friston and colleagues into a powerful framework based on Bayesian modeling [3], associating such concepts as perceptual inference [4], reinforcement learning [5] and optimal control [6]. In this framework, it has been proposed that the minimization of free energy -a concept coming from thermodynamics-could provide a general principle associating perception and action in interaction with the environment in a coherent predictive process [7, 8]. A number of recent neurophysiological studies confirm the accuracy of the predictive coding paradigm for analyzing sensory processing in the human brain (e.g. [9]).

Actually, predictive coding is a general methodological paradigm in information processing that consists in analyzing the local regularities in an input data stream in order to extract the predictable part of these input data. The information processing system can then focus on the difference between input data and their prediction. In a very general manner, whatever the processing system, there are two main advantages to processing the difference signal over directly processing the input signal. First, if the prediction is efficient, the difference signal is generally of (much) lower energy than the original signal, which leads to energy consumption saving in subsequent processes and resource saving for representing the signal with a given accuracy (e.g. bitrate saving in an audio or a video coder). In short, this reduces the “cost” of information processing. Second, there is a concentration of novelty / unpredictable information in the difference signal, which is exploitable for, e.g., the detection of new events. Because of these advantages, predictive coding has been largely exploited in technological applications, in particular in signal processing for telecommunications [10, 11].

### Predictions in speech neurocognitive processing

The present paper focuses on the concepts of predictability and prediction in the field of auditory processing, and also visual processing, for speech perception. Behavioral evidence for the existence of predictive processes in audition and speech perception actually appeared about 50 years ago, through such findings as forward masking in psychoacoustics [12] and selective adaptation in speech perception [13–15]. These two phenomena can be conceived as instances of predictability: neural detectors would adapt and decrease their response to the predictable input, while non-adapted neurons would be ready to detect novelty. The decrease of the neural response to predictable inputs in the auditory system has been largely displayed within animal unitary fibers [16]. Neural adaptation in the auditory nerve has an important role for the coding of speech phonetic cues [17–19].

A number of recent papers address the question of neural architectures likely to implement predictive coding for speech processing in the human brain. In the context of multiplex coding theories, [20] proposes that the predictions in time (“when” would something important happen) are based on a coupling between low-frequency oscillations driven by the syllabic rhythm in the “delta-theta” channel of neural firing (around 2-8 Hz) and mid-frequency regulation in the “beta” channel of neural firing (12-30 Hz) possibly related to the motor system. The “what” information would combine top-down predictions conveyed by the beta channel with analysis of the sensory input providing prediction errors to be conveyed to higher centers in a bottom-up process through the “gamma” channel (30-100 Hz).

### Acoustic, phonetic and linguistic sources of prediction in speech

Predictability of the speech input can be decomposed into various types of components: acoustic, phonetic and linguistic, respectively associated to different temporal scales that we call here short-, mid-, and long-term.

#### Short-term (acoustic level)

At the shortest temporal range, the acoustic structure of the speech signal provides a first source of predictions related to the fact that the speech signal is produced by a sound source (harmonic source from the vocal folds or noise source induced by a constriction inside the vocal tract) modulated by the vocal tract resonances. This results in regularities in the acoustic signal, captured by classical predictive coding techniques which have been intensively applied to speech signals. The vast majority of standardized predictive speech codecs used in telecommunications apply a so-called “short-term” prediction, i.e. prediction of a speech signal waveform sample from a linear combination of the preceding samples in the range of about 1 ms [21]. This is the basis of the famous linear predictive coding (LPC) technique and LPC family of speech coders. The predictor coefficients are calculated over individual short-time signal frames of a few tens of ms (typically 20-30 ms), every 10-20 ms.

#### Mid-term (phonetic level)

Predictions at a larger temporal scale are related to regularities in the structure of phonetic events in time. The dynamics of speech sounds are essentially organized into syllabic structures (typically at a time-scale around 100 to 200 ms) and include quasi-stationary regions (typically within vowels and fricatives or in the silent or voiced regions of plosive closure), phoneme-to-phoneme transitions within a limited series of possibilities in a given language (typically with a duration around 50 to 100 ms) and some larger coarticulation phenomena, typically based on anticipatory processes, that can reach several hundreds of milliseconds [22, 23]. In the field of speech coders, a few studies have applied some form of predictive coding on vectors of parameters encoding the speech information present in the short-term frame, using differential coding [24], recursive coding [25, 26], or Kalman filtering [27].^1^ Still, these approaches are limited to 1-step frame prediction, because of constraints on latency in telecommunications. Some other studies [28–34] have proposed “middle-term” speech coders, which aim at exploiting the speech signal redundancy and predictability over larger time spans, typically in the range of a few hundreds of ms. However, these methods actually implement a joint coding of several short-term frames (basically, by using trajectory models or projections), but do not apply any explicit prediction of a frame given past frames.

#### Long-term (linguistic level)

Finally, at an even larger temporal scale, long-term predictions are provided by the linguistic structure at the lexical, syntactic, semantic and pragmatic levels. The exploitation of this high-level linguistic structure is the heart of statistical language models (SLMs). SLMs are predictive models trained to estimate the posterior probability of observing a given word based on the other words already observed in the sequence (i.e. from past and/or future words). SLMs are used in automatic speech recognition (ASR) systems as a source of prior information for the acoustic-phonetic decoding process. However, to the best of our knowledge, they never concern the level of predicting a given piece of acoustic signal.

#### The role of vision

Importantly, the visual input can also bring some relevant information for phonetic predictions. As a matter of fact, pioneer studies such as [35, 36] showed that the visual component of an audiovisual speech input (e.g. “ba”) could result in decreasing the first negative peak N1 in the auditory event-related potential pattern in Electroencephalographic (EEG) data. Peak decrease has been related to the ability of the visual input to provide predictive cues likely to suppress the auditory response displayed in N1. The predictive aspect of visual speech information might be enhanced by the fact that there is often an advance of image on sound in natural speech (see [37]; though see a strong caveat in [38]). The potential predictive role of vision is supported by behavioral data showing that vision of the speaker’s face may indeed provide cues for auditory prediction, e.g. [39, 40]. Still, studies on audiovisual speech coders capable to exploit correlation between audio and visual speech are extremely sparse. We can mention [41], but again this study mostly focuses on a joint coding process, here of audio and visual vectors. A cascaded coding scheme was also proposed in this study (see also [42]), but even there, there is no explicit prediction and even less a measure of predictability (other than bitrate saving).

### Our contribution: modeling and assessing mid-term predictability in acoustic and audiovisual speech

The goal of the present study is to quantify what is really predictable online from the speech acoustic signal and the visual speech information (mostly lips movement). To this aim, we propose to use a series of computational models based on artificial (deep) neural networks which are trained to predict future acoustic features from past information. We focus on mid-term prediction, that is a prediction at the level of sequences of multiple consecutive short-term frames (i.e. from 25 ms to 250 ms in our experiments). As concerns the acoustic modality, we compare two different representations of the acoustic signal: a low-level raw power spectrum representation and a higher-level mel-frequency cepstral representation. As concerns lip movements, we use close-range video sequences of the speaker’s lips (visual region-of-interest). Depending on the representation and configuration (audio or audiovisual), different network architectures are used to learn spatio-temporal acoustic and visual patterns from data such as feed-forward neural networks and convolutional neural networks. Moreover, we implement these models using two configurations, a multispeaker configuration aiming at exploiting speech features shared among different speakers and a speaker-adapted configuration probing the possibility to transfer representations learned on a given pool of speaker to a new one. Importantly, this second configuration attempts to mimic the capability of a human brain to adapt its predictive processing to a new voice and new face.

The choice of a statistical framework based on deep learning was motivated by its ability to extract successive levels of increasingly meaningful abstractions from raw data in order to learn and perform complex (e.g. non-linear) mapping functions. By combining different types of generic layers (e.g. fully-connected, convolutional, recurrent, etc.) and training their parameters jointly from raw data without the need to hand-craft high-level discriminative features, deep learning provides a unified and coherent methodology to solve feature extraction, classification and regression tasks. Deep learning-based models have led to significant performance improvement in both acoustic speech processing (e.g. ASR [43], speech enhancement [44]), audiovisual speech processing (e.g. audiovisual and visual ASR [45–47]), articulatory-to-acoustic mapping [48] and more generally for tasks involving speech-related biosignals (see [49] for a review). Thus, deep neural networks are here considered as relevant candidates for modeling human behavior in specific processing tasks, in the present case speech predictive coding in the brain. Though substantially different from biological neural networks, artificial deep neural networks provide a computational solution to cognitive questions and may thus provide some insights on the nature of biological processes [50].

The proposed computational models of predictive speech coding enabled us to address the following questions:

- How much of the future speech sounds can be predicted from the present and the past ones?
- If the visual input (i.e. information on the speaker’s lip movements) is added to the acoustic input (i.e. the speech sound), how much gain can occur in prediction, and what temporal window of visual information is typically useful for upgrading auditory predictions?

We will first describe our corpora and methodology in more details. Then we will present the results of simulations that were carried out so as to attempt to answer the above questions, in the framework —and limitations–- of the available corpora and machine learning techniques exploited in this work.

## Material and Methods

### Data

#### Database

The publicly available NTCD-TIMIT database was used in this study [51]. NTCD-TIMIT contains audio and video recordings of 59 English speakers each uttering the same 98 sentences extracted from the TIMIT corpus [52] (i.e. 5,782 sentences in total). NTCD-TIMIT contains both clean and noisy versions of the audio material. In the present study, only the clean audio signals were used. As for the video material, NTCD-TIMIT provides a post-processed version of raw video sequences of the speaker’s face focusing on the region of interest (ROI) around the mouth. This includes cropping, rotation, and scaling of the extracted ROI so that the mouths of all speakers approximately lie to the same horizontal line and have the same width. Each ROI image is finally resized as a 67 × 67 pixels 8-bit grayscale image [51].

#### Data preprocessing

This section presents only the key steps of the data preprocessing. More details can be found in the Data processing section in *SI text*.

As concerns the audio recordings, a sliding window was used to segment each waveform into short-term acoustic frames. A classical frame length of 25 ms was used in our study (i.e. 400 samples at 16 kHz). Importantly, a frame shift of 25 ms was chosen in order to avoid any overlap between consecutive frames (i.e. the frame shift was set equal to the frame length). This aimed at preventing the introduction of an artificial correlation due to shared samples which could introduce some bias in the mid-term prediction (i.e. the prediction of a speech frame given the preceding ones).

Two approaches were then considered for further encoding the spectral content of each speech recording. In the first one, we computed the log-magnitude of the STFT spectrogram (on a dB scale), and re-scaled the resulting values to the range [0,-80] dB, for each sentence of the dataset.^2^ Each acoustic frame is here represented by a 257-dimensional vector containing the value of the power spectrum (in dB within the range [0,-80]) for 257 frequency bins linearly distributed in between 0 and 8 kHz. In the following we refer to this representation as the log-magnitude spectrogram.

In the second approach the short-term speech spectrum is converted into a set of 13-dimensional feature vector of so-called Mel-frequency cepstral coefficients (MFCC). MFCC coefficients are widely used in many fields such as ASR [53] and music information retrieval (e.g. classification of musical sound) [54]. Compared to using the raw STFT log-magnitude spectra as acoustic features, MFCC analysis can be seen as a high-level biologically-inspired process related to psychoacoustics (i.e. simulating the cochlear filtering). In our context, the use of MFCC coefficients as acoustic features may allow the computational model to focus only on the prediction task rather than on the extraction of discriminative features from the raw signal. Moreover, MFCC analysis leads to a more compact representation of the short-term speech spectrum (i.e. 13- vs. 257-dimensional feature vectors), which may be of significant interest in the context of statistical learning since it may limit the number of free parameters to estimate. All the above audio analysis procedures were performed using the *Librosa* Python open-source library [55].

As concerns the video sequences, a linear interpolation across successive images in the pixel domain was performed in order to adjust the video frame rate (originally 30 fps) to the analysis rate of the audio recordings (i.e. 40 Hz). Each 67×67 pixels frame was then resized to 32 × 32 pixels using linear interpolation. 8-bit integer pixel intensity values were divided by 255 in order to work with normalized values in the [0, 1] range. Video analysis was performed using the *openCV2* (Open Source Computer Vision) Python open-source library [56].

### Computational models of speech prediction from audio-only data

#### Task

Let us denote x_*t*_ a D_*x*_-dimensional (column) vector of spectral audio features, which can be either a 13-dimensional vector of MFCC coefficients or a 257-dimensional log-magnitude spectrum. The frame index *t* is an integer and corresponds to time *tH* where *H* = 25 ms is the frame spacing, see previous section. The predictive coding problem using past audio information can be formulated as computing: 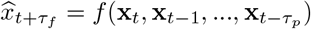, where 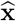 denotes an estimated (predicted) value of **x**, and *τ_f_* and *τ_p_* denote a time lag in the future and in the past, respectively (in number of frames). *f* is the non-linear predictive function modeled by the artificial neural networks described below.

#### Architectures

Two network architectures were used, depending on the considered representation of the speech spectrum. As concerns the log-magnitude spectrum, a convolutional neural network (CNN) [57] was used as the core of the proposed computational model, as shown in Fig. 1(a). A CNN is a powerful network architecture well-adapted to process 2D data for classification and regression. It can extract a set of increasingly meaningful representations along its successive layers. It is thus widely used in image and video processing, e.g. object detection [58], gesture recognition [59–62], or visual speech recognition [63] [47]. In the field of audio processing, CNNs have been applied to learn spatio-temporal regularities directly from sound spectrograms. Indeed, a time-frequency representation can be interpreted as a 2D image of the sound, and processed as such. With this approach, CNNs (in conjunction with other learning techniques such as recurrent neural networks) have shown very interesting performances in ASR [43] and audio events classification [64]. They are thus a natural choice for applying predictive regression over log-magnitude spectrograms. Note that, in the present study, the input image to the CNN consists of the portion of a speech spectrogram composed of frames *t* – *τ_p_* to *t*, i.e. a matrix [**x**_*t*_−*τ_p_* … **x***_t-1_* **x***_t_*], and eq-pred-1 can be reformulated as: 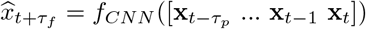. Technically, a CNN is a deep (multi-layer) neural network classically composed of one or several convolutional layers, pooling layers, fully-connected layers and one output layer. In a nutshell, a convolutional layer convolves an input 2D image with a set of so-called *local filters*, and then applies a non-linear transformation to the convolved image. The output is a set of so-called *feature maps*. Each feature map can be seen as the (non-linear) response of the input image to the corresponding local filter. One important concept in the convolutional layer is *weight sharing* which states that the parameters of each filter (called *weights*) remain the same whatever the position of the filter in the image. This allows the CNN to exploit spatial data correlation and build translation-invariant features. A *pooling layer* then downsamples each feature map in order to build a scale-invariant representation. For example a so-called *max-pooling* layer outputs the max value observed on sub-patches of a feature map. The convolutional + pooling process can be cascaded several times. A CNN generally ends up with a series of fully-connected layers which have the same function as in a standard feed-forward neural network: Each neuron of a given fully-connected layer performs a non-linear transformation of a weighted sum of its inputs (i.e. the outputs of the previous layer), except for the last (output) layer in the case of a regression task, where the output is directly the weighted sum. Note that in a CNN, the first fully-connected layer usually operates over a vectorized form of the downsampled feature maps provided by the last pooling layer.

**Figure 1.**
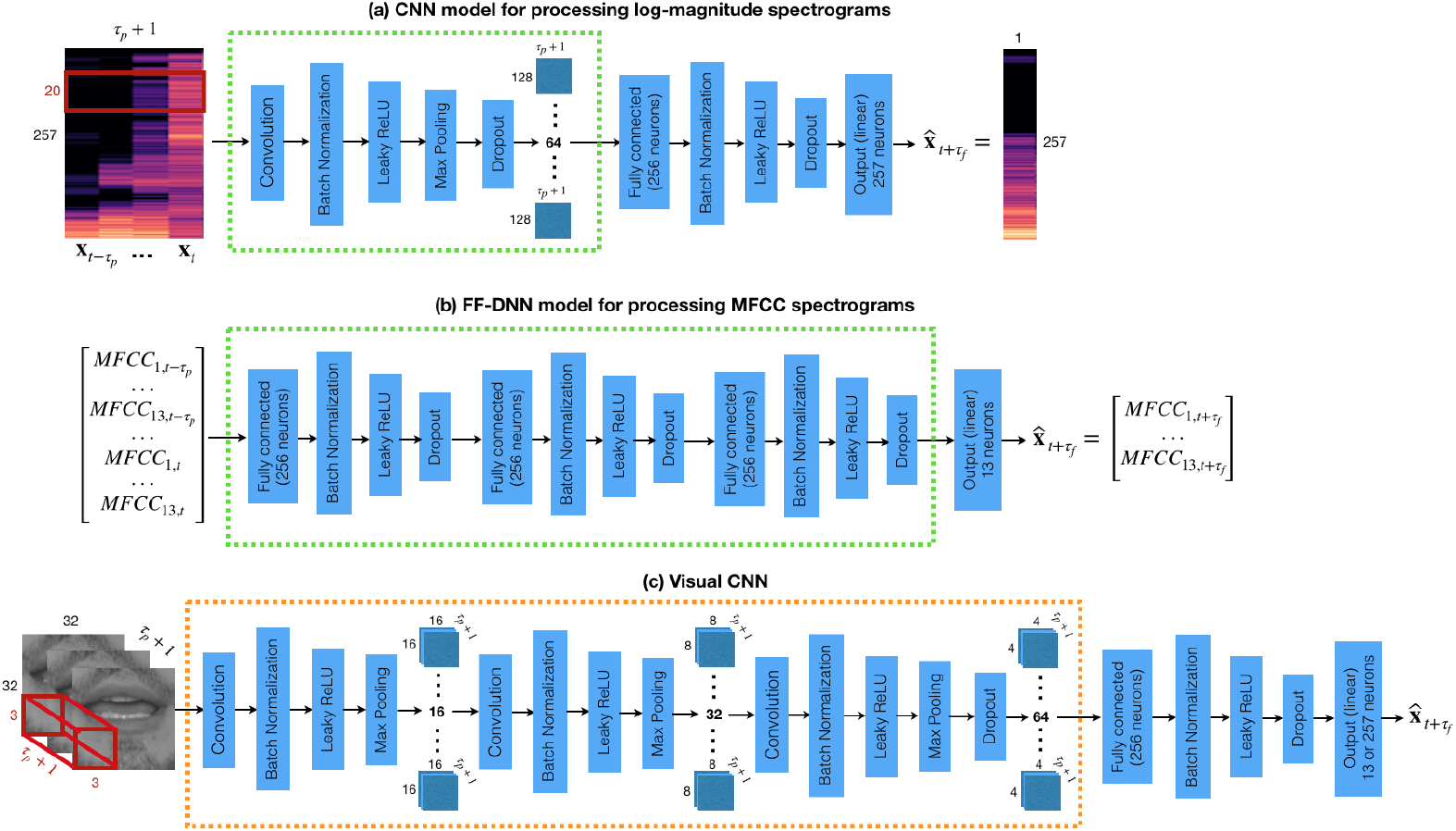
Selected architectures for the monomodal models and the cross-modal model. (a) CNN model for processing log-magnitude spectrograms; (b) FF-DNN model for processing MFCC spectrograms; (c) CNN model for predicting either a log-magnitude spectrum or an MFCC vector from a sequence of lip images. In (a) and (c) the red rectangle/cube represents the 2D/3D filters of the convolutive layer(s), respectively. The green/orange dotted rectangles indicate the subnetworks to be used in the audiovisual model (see Fig 2).

**Figure 2.**
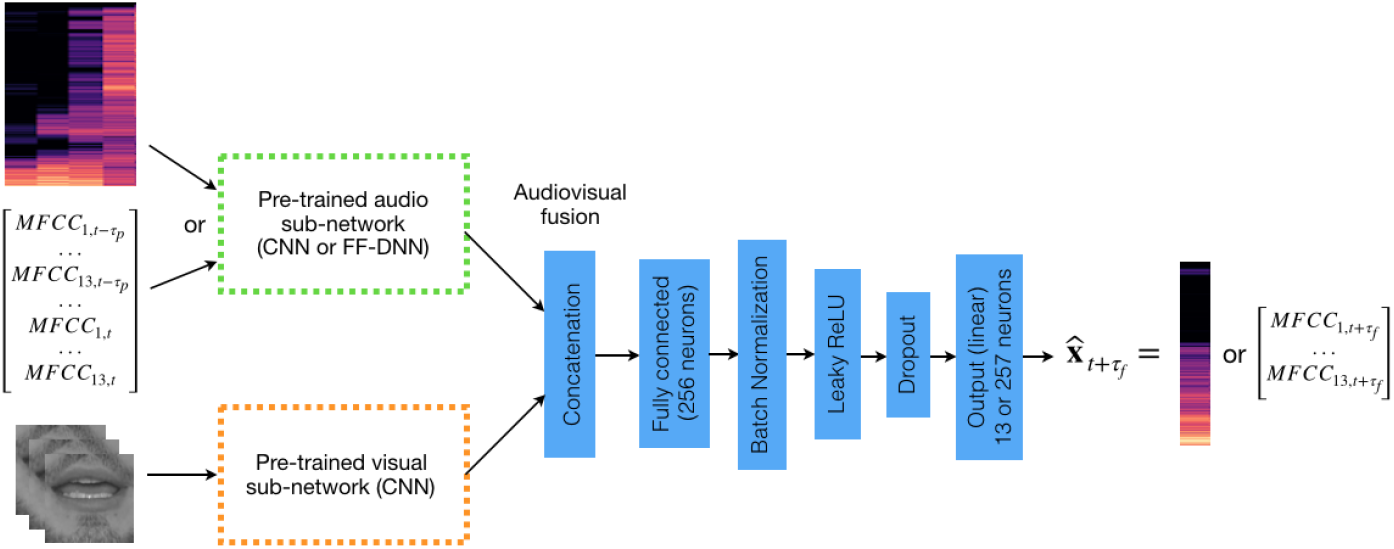
Selected architecture for the audiovisual model. Audio and visual pre-trained subnetworks from Fig 1 are merged using a 256-neuron fully-connected fusion layer. This architecture is valid for both log-magnitude spectrogram and MFCC spectrogram audio representations (the corresponding audio subnetwork is used).

As concern the MFCC spectrograms, the use of a CNN to learn regularities in the time-frequency domain is less meaningful than with log-magnitude spectrograms, since the DCT transformation involved in the MFCC calculation (see Data preprocessing section in SI-text) does not preserve locality in the extracted features. In other words, each MFCC coefficient is not related to a particular frequency range, and intra-frame correlation of MFCC coefficients is thus not necessarily concentrated on neighboring coefficients (actually, the DCT transform has the property to provide partially decorrelated coefficients). Therefore, to process the MFCC spectrograms, we rather use a “basic” feed-forward deep neural network (FF-DNN), as shown in Fig. 1(b). An FF-DNN is composed of a cascade of fully-connected layers, just as described above for the last layers of the CNN, including the last output layer which has a linear activation function. Note that, depending on the number of layers and number of neurons per layer, a FF-DNN is not necessarily less complex and less “powerful” than a CNN, in terms of number of free parameters (total number of weights) and modeling power, although the architecture is simpler. As opposed to the CNN, the input to a FF-DNN is technically a vector, not a 2D image. To process the portion of speech spectrogram composed of frames *t* − *τ_p_* to *t*, the latter has to be vectorized, i.e. concatenated into a single larger vector, and eq-pred-1 is here reformulated as: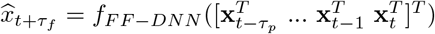, where ^T^ denotes vector transpose.

#### Model selection and training

All parameters (i.e. weights) of CNNs and FF-DNNs are learned from data, usually by stochastic gradient descent and backpropagation. Briefly, this consists in iterating the following process: (i) evaluating a loss function which measures the average discrepancy between the prediction of the network and the ground-truth value for a subset of the training data (called mini-batch), and (ii) calculating the gradient of this loss function with respect to all the network weights, starting from the output layer and back-propagating it through all the hidden layers, (iii) updating all weights using the gradient in order to decrease the loss function. This process is applied over all mini-batches of the training data, and repeated a certain number of times called epochs, until the loss function no more significantly evolves.

Importantly, as in many modeling studies based on deep learning, complex architectures such as FF-DNNs or CNNs require to set a large number of hyperparameters, mostly related to the sizing of the network, a process known as model selection. The procedure adopted in the present study for model selection and training is detailed in the Model selection and training section of the *SI Text*. The resulting models are the ones represented in Fig. 1(a) and 1(b).

### Computational models of speech prediction from both audio and visual data

#### Task

Let us denote **I**_t_ the lip image at frame *t* (in our case a 32×32 pixels grayscale image). The predictive coding problem using both audio and visual past information can be formulated as: 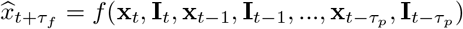.

##### Architectures

Integration of audio and visual speech information has been largely considered for automatic audiovisual speech recognition [45, 65] and also (though much less extensively) for other applications such as speech enhancement [66] and speech source separation [67]. Basically, the general principle is that integration can be processed at the input signal level (concatenation of the input data from each modality, aka early integration), at the output level (combination of the outputs obtained separately from each modality, aka *late integration*), or somewhere in between those extremes (after some separate processing of the inputs and before final calculation of the output, aka *mid-level integration*) [68]. Artificial neural networks provide an excellent framework for such multimodal integration, since it can be easily implemented with a fusion layer receiving the inputs from different streams and generating a corresponding output. Moreover, the fusion layer can be placed arbitrarily close to the input or to the output.

In the present study, we adopt the mid-level fusion strategy. Indeed, it enables to benefit from (i) the design and training of the audio (CNN or FF-DNN) network used for predictive coding based on audio only presented in the previous section, and (ii) the design and training of a visual CNN dedicated to process the lip images, since, as mentioned above, CNNs are efficient for feature extraction from images in classification and regression problems. We thus first designed and trained such a visual CNN, which architecture is represented in Fig. 1(c). Then, the convolutional + pooling layers of the visual CNN and either the convolutional + pooling layers of the log-magnitude spectrogram CNN or the fully-connected layers of the MFCC FF-DNN are selected. These subnetworks are merged using a fully-connected fusion layer, which is followed by other usual layers. The whole resulting network regressing audio and visual data into audio data is represented in Fig. 2. This audiovisual model was anew trained with the audiovisual training data. Again, further details about model selection and training are given in the corresponding section in *SI-Text*.

#### Multispeaker and speaker-adapted training

The training and evaluation of the models was done following two different paradigms and associated configurations. In the first one, referred to as *multispeaker* training, we aimed at assessing the generic nature of the predictions. For this aim, the entire NTCD-TIMIT corpus (59 speakers × 98 sentences = 5,782 audiovisual sequences) was randomly partitioned into 3,099 sentences for training 774 sentences for validation, and 1,909 sentences for test. Here, training, validation and test sets may share data from the same speaker, but of course for different sentences.

In the second configuration, referred to as *speaker-adapted training*, we assessed the ability of a model trained on a pool of speakers to process data from a new speaker, after being adapted with only a limited amount of data. The detail of this configuration is the following: (i) For each of the 59 speakers, the 98 sentences were divided into 66 for training/validation and 32 for test. The partition was drawn randomly but the same partition was applied to all speakers (so that a given sentence in a speaker-specific test set is never used for training the initial multispeaker model). (ii) A multispeaker model was trained using a subset of the training data comprising 42 speakers, randomly chosen among the 59 speakers (hence 66 ×42 = 2, 772 sentences). (iii) This multispeaker model was then fine-tuned (i.e. further trained starting from the output of step (ii)) separately on each speaker-specific training/validation subset (i.e. 66 sentences), for each of the remaining 17 speakers not used in step (ii). Performance was then averaged over the 17 corresponding speaker-specific test subsets (hence over 32×17 = 544 sentences).

### Metrics

Two metrics were used to assess the prediction performance of the different models. First, we used the mean squared error (MSE) between the predicted audio vector and the corresponding ground-truth audio vector. This MSE was also used as loss function to train the different models. The second metric is the weighted explained variance (EV) regression score, evaluating the proportion to which the predicted coefficients account for the variation of the actual ones. These two metrics are strongly related to a key metric of the predictive coding theory, which is the prediction gain [10, 21], which is also discussed in the results. The corresponding equations for these metrics are detailed in the Metrics section of *SI-Text*.

## Results and discussion

The prediction performances of the models based on log-magnitude spectrogram and MFCC spectrogram are presented in Fig. 4 and Fig. 3 respectively.

**Figure 3.**
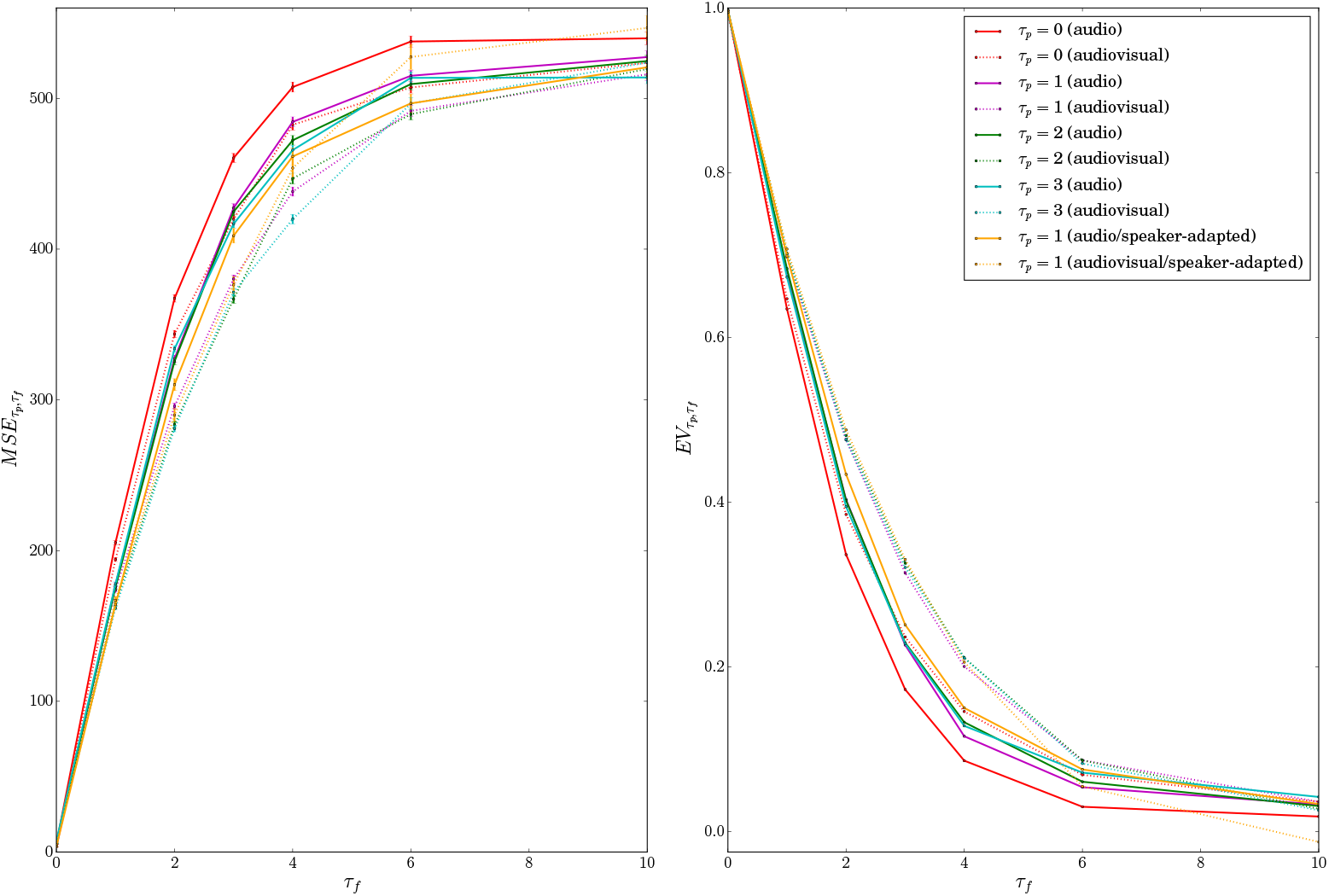
Prediction performance of the audio and audiovisual models using the MFCC spectrogram as audio representation in terms of mean square error (left) with 95% confidence interval (represented by the error bars) and explained variance regression score (right). All models were trained using the multispeaker configuration, except when indicated (where training was done using the speaker-adapted configuration).

**Figure 4.**
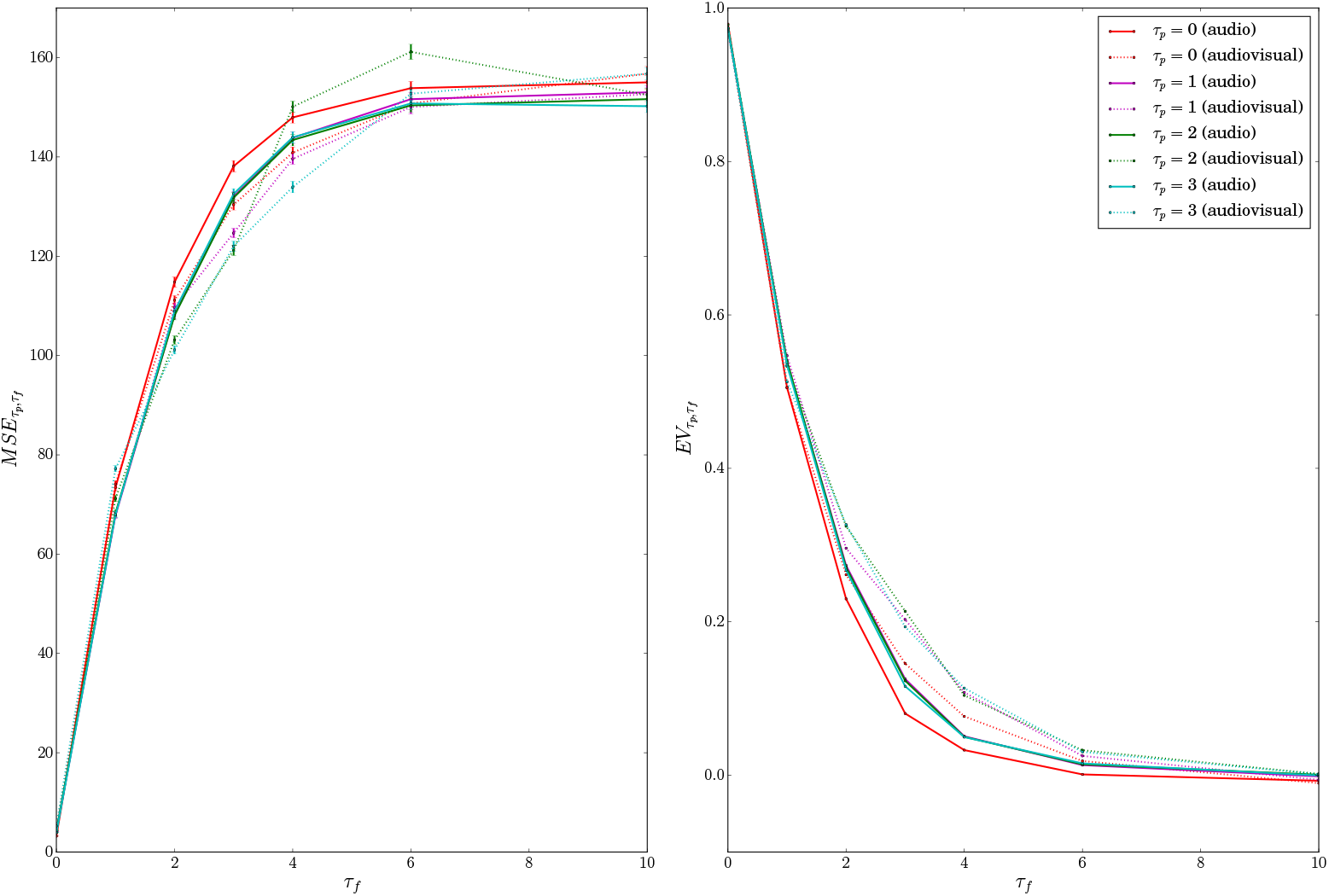
Prediction performance of the audio and audiovisual models using the log-magnitude spectrogram as audio representation in terms of mean square error (left) with 95% confidence interval (represented by the error bars) and explained variance regression score (right). All models were trained using the multispeaker configuration.

### Global tendencies

The results show that it is indeed possible to predict, to a certain extent, the spectral information in the acoustic speech signal in a temporal range of 100 ms following the current frame (*τ_f_* ≤ 4). As expected the accuracy of such predictions decreases rapidly when the temporal horizon *τ_f_* increases. This evolution more or less follows a logarithmic shape for 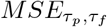 and an exponential decay toward almost 0 for 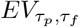.

At 25 ms (*τ_f_* = 1), the weighted explained variance is about 0.5 for the audio-only model exploiting (and predicting) the log-magnitude spectrogram (e.g. for *τ_p_* = 3, *E V* _3,1_ = 0.53) and about 0.65 for the one based on the MFCC spectrogram (e.g. for *τ_p_* = 3, *EV*_3,1_ = 0.67; compare cyan solid lines in Fig. 4 and Fig. 3). This corresponds to an average predictive coding gain *G_τ__p_,_τ__f_* around 2 and 2.8, respectively. These values mean that the power of the error signal (input minus predicted) to transmit by neural processes is reduced by a factor between 2 and 3 compared to the original input, which provides some quantification of the amount of biological energy that the system might gain in exploiting a predictive process.

At 50 ms (*τ_f_* = 2), depending on the audio representation, the weighted explained variance is about 0.4 (e.g. for *τ_p_* = 2, *EV_2,2_* = 0.4 for the audio-only model based on MFCC spectrogram) which corresponds to an average prediction gain around 1. 7. For the two audio-only models, prediction becomes almost random at 100 ms and completely vanish at 150ms (e.g. *EV*_2_,4= 0.13 and *EV*_2_,6= 0.07 for the audio-only model based on MFCC spectrogram). This provides a rather accurate estimation of the temporal window in which acoustical predictions are available, typically around half a syllable in time.

### Impact of audio representation

The use of MFCC spectrogram as audio representation systematically leads to better prediction than the use of STFT log-magnitude spectrogram (see Fig. 4 *vs*. Fig. 3). For instance, *EV*_2,2_ = 0.4 for the audio-only model based on MFCC spectrogram whereas *EV*_2,2_ = 0.28 for the same model but based on log-magnitude spectrogram. To interpret the differences, let us recall that, compared to the log-magnitude spectrogram, the MFCC spectrogram is much more *compact* (13 vs. 257 coefficients to predict) and more *high-level* since it focuses on the low frequency range which conveys most of the phonetic information (while in the log-magnitude spectrum each coefficient, i.e. each frequency bin, has an equal weight on the prediction accuracy). These two aspects may lead to more robust and more efficient predictive models.

### Impact of past information

Another aspect of acoustic prediction concerns the role of the temporal context. Unsurprisingly, adding one context frame to the current one provides significant improvement in the prediction of the next frame (e.g. comparing results for *τ_p_* = 0 and 1 in Fig. 3). Such past information may enable the model to evaluate speech trajectories and extract relevant information on the current dynamics, related e.g. to formant transitions, known to be crucial in speech perception. However, adding a second frame of past context (*τ_p_* = 2) does almost not improve prediction (only for *τ_f_* ≥ 4 for which the prediction accuracy is already low and only for experiments based on MFCC spectrogram). Adding a third one (*τ_p_* = 3) just maintains most performances. While being related to a different task, such results may be compared to classical ones in automatic speech recognition, where adding first and second derivatives of the spectral parameters is classically considered as the optimal choice for reaching the best performance.

### Prediction accuracy per class of speech sound

A fine-grained analysis of the prediction accuracy for five major classes of speech sounds is presented in Fig. 5. Interestingly, the prediction accuracy at *τ_f_* = 1 is in a comparable range for vowels, fricatives, nasals and semivowels but is significantly lower for plosive sounds (i.e. significantly larger *M S E*). This may be explained by the difficulty to predict the precise timing of the occlusion release within the plosive closure, and to predict the shape of the corresponding short-term spectrum from pre-release signal. This pattern is also visible but decreased for predictions at 50 ms and 75 ms (i.e. *τ_f_* = 2 and *τ_f_* = 3) probably due to a ceiling effect of the prediction accuracy at these temporal horizons.

**Figure 5.**
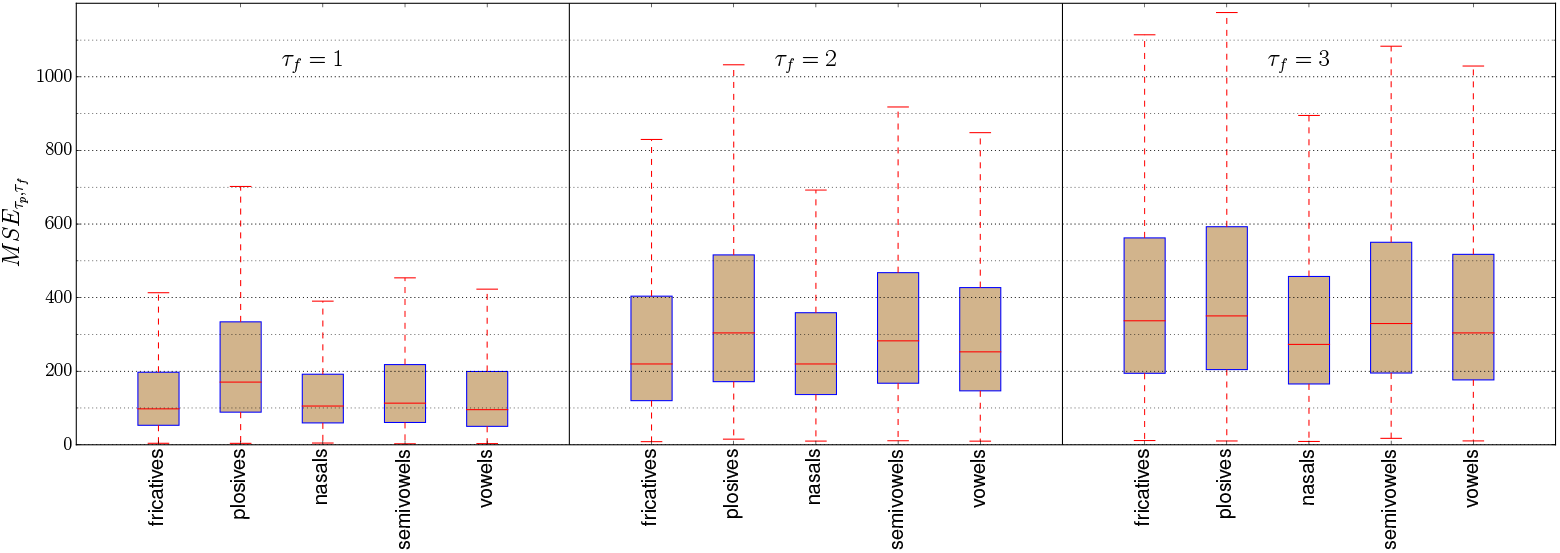
Prediction accuracy per class of speech sound in terms of *M S E*_*τ_p_*,*τ_f_*_ for *τ_f_* ∈ [1, 2, 3] (i.e. 25,50,75 ms) using an audio-only predictive model based on MFCC spectrogram and trained using the multispeaker configuration with **τ_p_** = 3.

### Impact of training paradigm

Now we discuss the results obtained with speaker-adapted training, and compare them to multispeaker training. We recall that speaker-adapted training aims at probing the ability to transfer some representation learned from a pool of speakers to a new one, given only a small amount of adaptation data (audio-only or audiovisual). The results are presented in Fig. 3 (see orange solid lines for audio-only models and orange dashed lines for audiovisual models; for purpose of brevity and clarity, we reported only the results obtained when using the MFCC spectrogram as audio representation and for τp = 1). Interestingly, for prediction up to 125 ms (up to *τ_f_* = 6), the accuracy of the speaker-adapted audio-only model is significantly better than the same multispeaker model (e.g. *M S E*_1,5_ = 461 with a speaker-adapted model and *M S E*_1,5_ = 484 with a multispeaker model, with 95% confidence interval equal to 5). This shows the possibility for a predictive model of speech to adapt to any new arbitrary speaker, at least for mid-term prediction.

### Impact of visual input

First, we present in Fig. 6 (left) the performance of visual-only models (i.e. predictive models of acoustic speech that rely *only* on visual information). As expected, the information provided by the visual modality is real but limited. For example, the best performance obtained at *τ_f_* = 0 is *EV*_3,0_ = 0.37 only. This result can be put in perspective with respect to the literature on automatic lip-reading (also known as visual speech recognition) where a typical performance of a visuo-phonetic decoder which does not exploit any high-level linguistic knowledge (via SLMs) is between 30% and 40% (i.e. 60-70% phone error rate). Similarly to the audio-only models, adding past context frames to the current one provides significant improvement in the prediction accuracy. Most of this improvement is observed when considering past information at 25 ms (i.e. τp = 1) but adding one more past context frame (i.e. *τ_p_* = 2) leads to the best performance. Interestingly, the performance of visual-only models decreases relatively slowly, e.g. at 50 ms *EV*_2,2_ = 0.32 and at 75 ms *EV*_2,3_ = 0.25. How*EV*er, on average, visual modality does not seem to convey useful information above 100ms (e.g. at 150ms, *EV*2,6 = 0.06). These results may contribute to the debate in the neuroscience literature on the fact that the lip movements could be in advance on the sound, because of anticipatory processes in speech production (see, e.g., [37, 69]). As mentioned above, the prediction of the spectral parameters from the lip information is maximal for the frame synchronous with the current input lip image (i.e. *τ_f_* = 0) (or for the next image only for the configuration (τp = 2, *τ_f_* = 1)) and then decreases smoothly with time. This is not in agreement with a stable advance of lips on sound.

**Figure 6.**
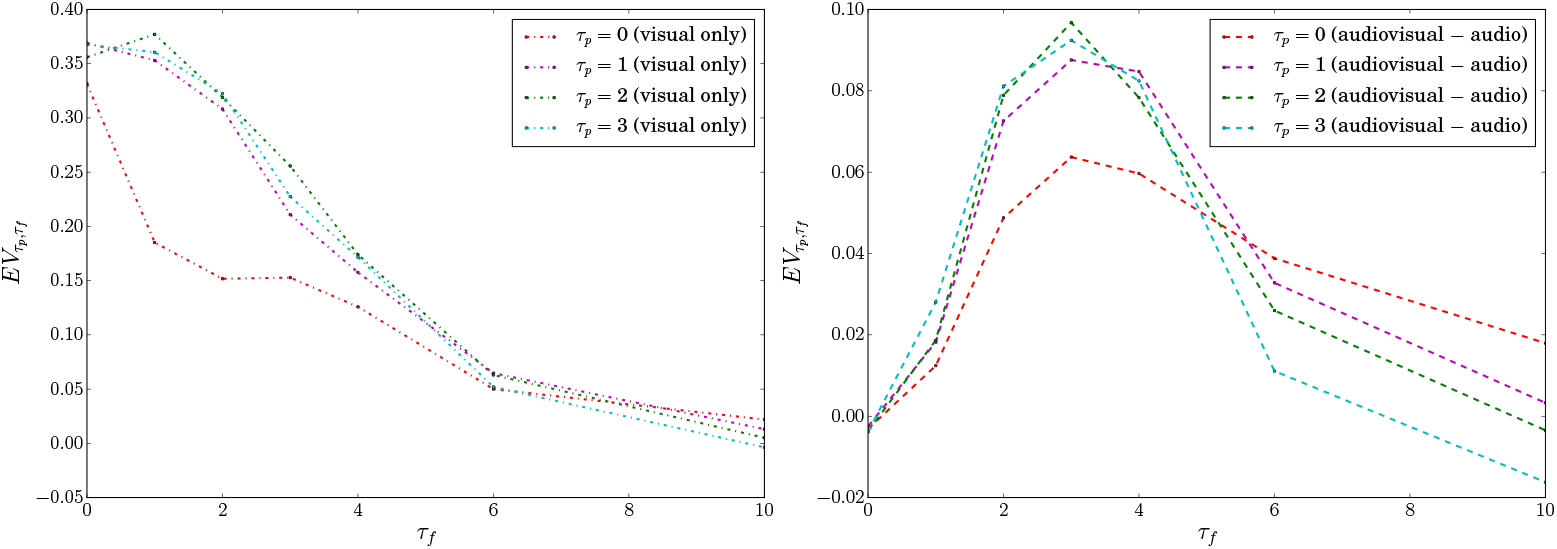
Explained variance obtained when considering only the visual modality (left) and difference of explained variance between audio and audiovisual models (right) considering the MFCC spectrogram as audio representation and the multispeaker training configuration.

As illustrated in Fig. 4 and Fig. 3, combining audio with visual information improves the prediction (compare solid lines with dashed lines). The gain is small but real, increasing weighted explained variance by 0.01 to 0.1. It is more important for the multispeaker training configuration than for the speaker-adapted one (with better performance of the speaker-adapted models observed only for prediction below 100 ms and a bit surprisingly, lower performances observed for larger temporal ranges). This may r*EV*eal a difficulty to transfer internal representations related to the visual input to a new speaker with only a limited amount of adaptation data (adapting the model to a new morphology and to speaker-specific articulatory strategies).

In order to better illustrate the dynamics of the gain brought by the visual input, we displayed in Fig. 6 (right) the difference of explained variance between audiovisual models and audio-only models (for the multispeaker training configuration).

Importantly, results show a peak in the gain provided by the visual input for a delay of 75 ms (τp = 3): this is where the prediction error is maximally decreased and the weighted explained variance is maximally increased by the visual input. Therefore, even if there is no systematic lead of lips on sounds, there is a range of asynchronies between 50 ms and 100 ms where the gain due to the visual input is rather stable.

### Qualitative *EV*aluation of a predicted spectrogram

All the above-presented quantitative results were averaged over many test sentences and speakers. Here, we finally discuss from a qualitative point of view the accuracy of the predicted spectral content at the utterance level. Examples of predicted log-magnitude spectrogram at 50 ms (i.e. *τ_f_* = 2) using either an audio-only, a visual-only and an audiovisual predictive model are shown in Fig. 7. As concerns the audio-only model (see blue in plot (g)), peaks in prediction errors are mainly observed either at the vowel onset of consonant-vowel sequences (e.g. [s-uw], [iy-aa], [d-aa]) or at the onset of the consonant of vowel-consonant sequence (e.g. [er-k], [er-t]) for which the precise initiation of the trajectory after a period of relative stability is hardly predictable. As concerns the audiovisual model, the average gain is accompanied by a large range of variations, leading to fluctuations between large gains and large losses provided by lip movements.

**Figure 7.**
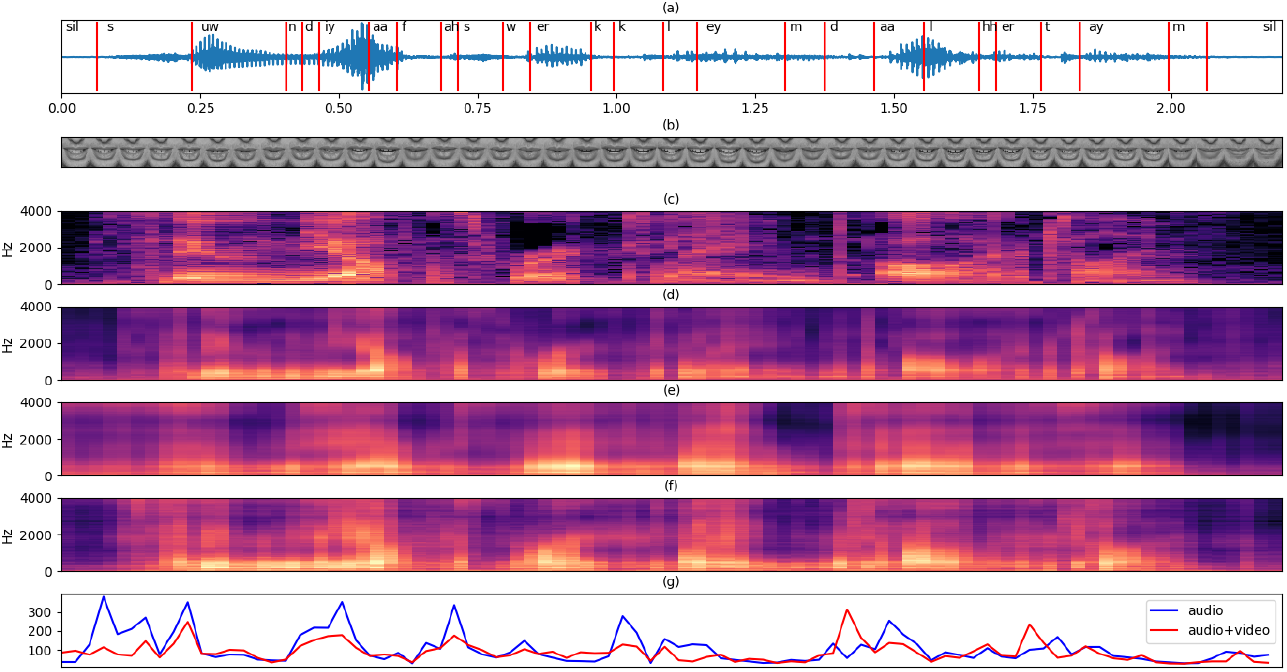
Examples of reconstructed log-magnitude spectrogram for *τ_p_* = 3 and *τ_f_* = 2 using an audio-only, a visual-only and an audiovisual predictive model. (a) original audio waveform with phonetic segmentation (with TIMIT phonetic labels) for sentence *sil100* recorded by speaker *57M* of NTCD-TIMIT corpus, (b) corresponding lip image stream (downsampled to 15fps to improve figure readability), (c) original log-magnitude spectrogram (display only between 0 and 4kHz for readability), (d) log-magnitude spectrogram estimated from (past) audio-only observations (M S E_*τ_p_*,*τ_f_*_ = 106.8,*E V*_*τ_p_*,*τ_f_*_ = 0.3), (e) log-magnitude spectrogram estimated from (past) visual-only observations (*M S E*_*τ_p_*,*τ_f_*_ = 116,*EV*_*τ_p_*,*τ_f_*_ = 0.2), (f) log-magnitude spectrogram estimated from both (past) audio and visual observations (*M S E*_*τ_p_*,*τ_f_*_ = 88,*EV*_*τ_p_*,*τ_f_*_ = 0.4) (g) Temporal evolution of *M S E*_*τ_p_*,*τ_f_*_ over the utterance for both audio and audiovisual predictive models.

A substantial gain from the visual input may occur when the speaker produces preparatory lip gestures before beginning to speak as in the [s] onset after the silence at the beginning of the utterance. Another source of gain could be related to a co-articulation effect, when lips anticipate the upcoming vowel as during the first [s] in the utterance where the protrusion gesture starts before the onset of the [uw]. Conversely, cases of error increase due to the visual input concern occurrences of non-visible tongue gestures, e.g. intensity decrease in the vowel [er] due to tongue closure in the following [t] around 1.75 s that is detected in the auditory stream but not in the visual stream.

### A database of prediction errors for future neurocognitive experiments

A number of recent neurophysiological experiments have tested the existence and characteristics of predictive patterns in the audio and audiovisual responses to speech in the human brain (see, e.g., [36, 70–72] and a recent theoretical r*EV*iew of predictive processes [9]). Importantly, these experiments lack a ground truth basis on the natural predictive structure of audio and audiovisual speech, which can lead to misinterpretations or over-generalizations of observed patterns [38]. The present study could provide an interesting basis for future studies, providing a quantitative knowledge on the amount of “predictability” available in the physical signals considered in the simulations. The source code used to format the data, train and *EV*aluate both audio and audiovisual predictive models as well as all experimental data and results have been made publicly available (DOI: 10.5281/zenodo.1487974). We beli*EV*e that such results could be of interest for future neurophysiological experiments aiming at testing neural predictions in speech processing in the human brain.

## Conclusion

In the general framework of predictive coding in the human brain, the present study aimed at quantifying what is really predictable online from the speech acoustic signal and the visual speech information (mostly lip movements). We proposed a set of computational models based on artificial (deep) neural networks which are trained to predict future audio observations from 25 ms to 250 ms from past audio or audiovisual observations. The deep learning approach enabled us to process (almost) raw audio and visual data while minimizing the amount of prior information (and limit the possible biases brought by hand-crafted features). Model training and *EV*aluation were performed on a large multi-speaker database, publicly available. The key results of the present study are:

- It is possible to predict the spectral information in the acoustic speech signal in a temporal range of 100 ms. At 25 ms, prediction enables to reduce the power of the error signal – relative to prediction – to transmit by neural processes by a factor 2.5. But the accuracy of the prediction decreases rapidly (e.g. with average prediction gain around 1.5 at 50ms, 1.25 at 75ms and almost 1, i.e. no gain, at around 100 ms). This tendency is robust across audio representations (STFT log-magnitude vs. MFCC spectrogram).
- The information provided by the visual modality is real but limited. Prediction accuracy of the predictive model based on visual-only information does not *EV*idence a stable advance of lips on sound (as sometimes stated in the literature). Maximum average gain provided by the visual input in addition to the audio input (up to about +0.1 of explained variance) is obtained for a prediction at 75 ms.
- Best prediction accuracy is obtained when considering 50 ms of past context, for both audio and audiovisual models.
- Plosives are more difficult to predict than other types of speech sound.
- It is possible to adapt a (audio or audiovisual) predictive model of speech from a pool of speakers to a new speaker, given a small amount of adaptation data, without significant performance loss.

Finally, it is of importance to mention what misses in the present study, that is the linguistic levels, from words to sentences, which of course contain a large potential for further prediction. It is well known that language models provide a range of predictions for phonemes and words, that are used for instance in automatic speech recognition systems. Our choice here was to assess exclusively the sensory (audio and audiovisual) level independently of high-level linguistic predictions. Importantly, linguistic predictions require a large context and may induce large delays in prediction. Future work will focus on the integration of this linguistic level in a more complete neurocognitive architecture for speech predictive coding.

## Acknowledgments

This work has been supported by the European Research Council under the European Community Seventh Framework Programme (FP7/2007-2013 Grant Agreement no. 339152, Speech Unit(e)s.)

## Supplementary Information (*SI text*)

### Data preprocessing

Each audio recording (16-kHz, 16-bit) was first cropped in order to reduce the amount of silence before and after each uttered sentence. Temporal boundaries of silence portions were extracted from the phonetic alignment file provided with the NTCD-TIMIT corpus (obtained by an automatic HMM-based force-alignment procedure). In order to take into account anticipatory lip gestures, a safe-margin of 150 ms of silence was kept intact before and after each recorded sentence.

The discrete Fourier transform (DFT) was applied on each frame to represent its spectral content (we recall that each frame is 25 ms length and that there is no overlap between two consecutive frames in order to pr*EV*ent the introduction of an artificial correlation between them). The overall process is referred to as the Short-Term Fourier Transform (STFT) analysis and the resulting signal representation is the STFT (complex-valued) spectrogram. This is a very common time-frequency representation in the speech/audio processing literature, although it is quite rarely applied without overlap between frames, as is done here, because usual STFT analysis with overlap deliberately intends to introduce some correlation in between adjacent frames to obtain a smooth spectrogram in the time dimension. In the present study, a 512-point Fast Fourier Transform (FFT) was used to calculate each DFT (each 400-sample short-term frame was zero-padded with 112 zeros, and was then applied a Hanning analysis window). Only the 257 first coefficients in the frequency dimension, corresponding to positive frequencies, are retained.

In the second approach, called Mel-frequency cepstral analysis, subband integration of the log power spectrum was processed using a set of 40 triangular filters equally spaced on a nonlinear Mel frequency scale which aims at approximating the human auditory system’s response. The resulting 40-dimensional Mel-frequency log-spectrum was then converted into a 13-dimensional vector of so-called MFCC coefficients, using the discrete cosine transform (DCT). The resulting representation for a complete utterance (a sequence of frames) is referred to as the MFCC spectrogram. All the above audio analysis procedures were performed using the Librosa Python open-source library [55] (release 0.6.0).

### Model selection and training

As in many modeling studies based on deep learning, complex architectures such as FF-DNNs or CNNs require to set a large number of hyperparameters, mostly related to the sizing of the network, a process known as model selection. An extensive search for the optimal combination for these hyperparameters is out of range. Therefore, we optimized only some of them on a hold-out dataset containing the first 5 speakers among the 59 ones available in NTCD-TIMIT. 70% of this hold-out dataset was used for training the models in the model selection phase, and 30% was used for test.

As concerns the CNN model for processing audio-only data, we tested 1 or 2 groups of convolutional/pooling layers. For the convolution layer (and for each group), we tested different numbers of 2D filters *N_f_* ∈ {16, 32, 64, 100}, with different sizes along the frequency dimension *K_f_* ∈ {3, 5, 10, 20}. The filter size along the temporal dimension was fixed to *τ_p_* + 1, i.e. the maximum size adjusted to the considered temporal span of the inputs [t,t - *τ_p_*]. The pooling factor was fixed to 2×1 (i.e. the pooling was performed across the frequency axis within 2 consecutive frequency bins). We tested different numbers of fully-connected layers *N_FC_* ∈ {1, 2} with either 256 or 512 neurons each.

As concerns the CNN model processing video-only data (used to initialize theaudiovisual model), we tested 1, 2 or 3 groups of convolutional/pooling layers. As often done in computer vision tasks involving CNNs (e.g. [63]), the number of filters was incremented in each layer, e.g. 16 for the first layer, 32 for the second, 64 for the third. Filters of size 3×3, 5×5, 10×10 were tested. The pooling factor was fixed to 2×2. As for the audio CNN, we tested different numbers of fully-connected layers NFC ∈ {1, 2} with either 256 or 512 neurons each.

As concerns the FF-DNN used to process MFCC spectrograms, we tested combinations of 1, 2, 3 and 4 hidden layers with either 128, 256, or 512 neurons each. Finally, for the audiovisual model (i.e. the one jointly processing audio and visual data to predict audio), we used the subnetworks of the selected audio and visual networks, and we only varied the number of fully-connected layers *N_FC_* ∈ {1,2, 3}, with either 256 or 512 neurons each.

For all models, we tested the following activation functions: tanh, ReLU, and Leaky ReLU which is defined as *f* (*x*) = *αx* for *x* < 0 and *f* (*x*) = *x* for *x* ≥ 0 (with *α* = 0.03 in our experiments).

All models were trained using the Adam optimizer (a popular variant of the stochastic gradient descent) [73], on mini-batches of 256 observations. The mean squared error (M SE) was used as loss function. Three strategies were combined to pr*EV*ent model overfitting and accelerate training convergence: (i) early stopping which consists in monitoring the loss function on a validation dataset (i.e. hold-out dataset, 20% of the training set in our case) and stopping the training as soon as its value stops decreasing after a given number of epochs (set to 10 in this study); (ii) batch normalization which consists in applying a transformation so that the inputs to each layer have zero mean and unit variance [74]; (iii) dropout which consists in not updating a random fraction of neurons in a given layer during training. Technical implementation of all models was performed using the Keras open-source library [75] (release 2.1.3). All models were trained using GPU-based acceleration.

The selected audio CNN processing log-magnitude spectrograms is the one with a single group of convolution+pooling layers, with 64 filters of size 20× (*τ_p_* + 1), and a single 256-neuron fully-connected layer. The selected visual CNN processing lip images has 3 groups of convolution+pooling layers, with 16, 32 and 64 filters of size 3× 3× (*τ_p_* + 1), and a single 256-neuron fully-connected layer. The selected audio FF-DNN processing MFCC spectrograms has 3 groups of 256-neuron fully-connected layers. Finally, the audiovisual model merges the subnetworks from the selected audio and visual models, using a 256-neuron fully-connected fusion layer. These resulting models are the ones represented in Fig. 1 and Fig. 2.

## Metrics

Two metrics were used to assess the prediction performance of the different models. First, we used the mean squared error (*M S E*) between the predicted audio vector and the corresponding ground-truth audio vector. This *M S E* was also used as loss function to train the different models. For each pair (*τ_p_*, *τ_f_*) of “past context lag / prediction lag”, the *M S E* is first defined per frequency bin or MFCC coefficient (indexed by d) and per test sentence (indexed by *k*) as:

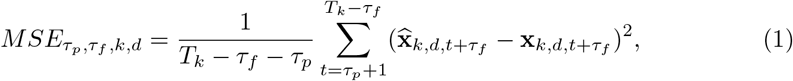

where *T_k_* is the number of audio vectors (i.e. the number of acoustic short-term frames) in sentence *k*, and 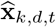 and **x***_k_,_d_,_t_* are the *d*-th entry of respectively the predicted and ground-truth vectors at frame t for the k-th sentence. Then it is averaged across frequency bins and across sentences:

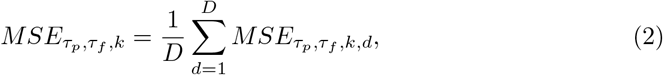

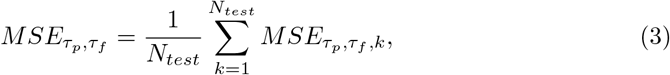

where *D* = 13 or *D* = 257 depending on the audio representation, and *N_test_* is the number of test sentences (i.e. 1,909 for the speaker-independent configuration and 544 for the speaker-adapted configuration). Assuming a Gaussian distribution of the errors, a 95% confidence interval of *M S E_τ_p_,τ_f__* is defined as:

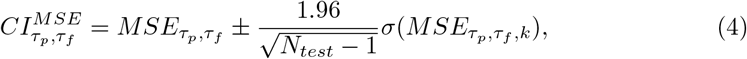

where *σ*(.) denotes the empirical standard deviation evaluated over the set of test sentences.

The second metric is the weighted explained variance (EV) regression score defined as:

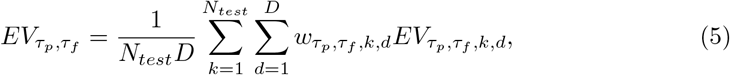

with

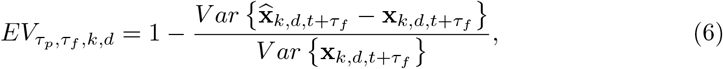

and

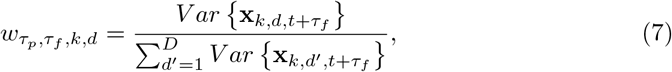

where *V ar* denotes the empirical variance evaluated along the time dimension (the different frames of a sentence). For each sentence, the contribution of the d-th frequency bin to the explained variance is weighted by the relative average power of the signal in that frequency bin. This avoids a bin with very low power to yield a very large (negative) *EV* value for that frequency bin, which would pollute the average *EV* value. In contrast, the M SE is not weighted. The weighted *EV* is lower than 1, with a value close to 1 indicating a strong correlation between predicted and ground truth data, and a lower value (potentially negative) indicating a poorer correlation.

Finally, it can be noted that both M SE and *EV* are strongly related to a key metric of the predictive coding theory, which is the prediction gain [10, 21]. Indeed, the latter is defined as the ratio of the ground truth signal power and the prediction error power (i.e the M SE):

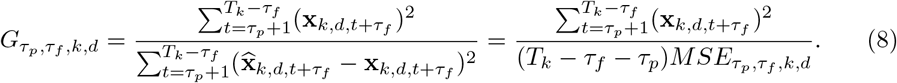

The lower the M SE the higher the prediction gain. Moreover, in the case where both ground truth signal and predicted signal are zero-mean, we have:

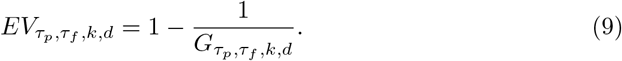

Therefore, the higher the prediction gain, the closer to 1 the expected variance. Those relations are given here for each frequency bin and each sentence. Depending on howaveraging across frequency and sentence is performed, they can become more intricate after averaging. In the present study, we define an average prediction gain such as:

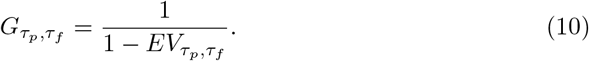

## Footnotes

1 So, when the short-term vector is composed of LPC coefficients, we thus have two chained levels of prediction: inter-frame prediction of an intra-frame predictor.

2 More precisely, the maximum value over each sentence was set to 0 dB and all values below −80 dB were set to −80 dB.

